# A mouse model of necrotic biliary pancreatitis induced by combining gallstone formation and ligation of the biliary-pancreatic duct

**DOI:** 10.1101/158915

**Authors:** Zuobiao Yuan, Junyuan Zheng, Zhu Mei, Guoyong Hu, Yue Zeng

## Abstract

**Objective:** The aim of the present investigation is to develop a mouse model of biliary pancreatitis with characteristics of both gallstone formation and pancreatitis, mimicking the human etiology and pathphysiological character.

**Design:** Male C57BL/6 mice were fed with chow, high fat/cholesterol and lithogenic diet for 12 weeks respectively. Laparotomy was done followed by ligation of pancreatic duct (PD), bile duct and pancreatic duct (BPD), or sham operation.

**Results:** Little or no evidence of pancreatitis was observed in PD group of mice fed with chow or high fat/cholesterol diet, or in the tail of pancreata removed from animals fed with lithogenic diet. In the head of pancreas, pancreas damage was dramatically more severe in the lithogenic group. When bile reflux was blocked by BPD, pancreas damage markedly reduced to level of chow diet group. The lithogenic diet group also developed significantly more severe multi organ dysfunction syndrome (MODS) in the lung, kidney and liver. The severity of pancreatitis is associated with persistent high bile level of cholesterol and bile acid after obstruction of the biliary-pancreatic duct. Cholesterol crystal aggravated injury of pancreatic acinar cells caused by taurocholate. After obstruction of the biliary-pancreatic duct, in the lithogenic diet group, liver Abcg8 and Cyp7a1 was up-regulated, compared to the control group.

**Conclusion:** We developed a mouse model of severe biliary pancreatitis in both local pancreas damage and MODS. This model provides a sound explanation for the Opie theory dilemma and a potential therapeutical direction in clinical practice as well.

**Summary statement:** A biliary pancreatitis has characters of both gallstone and pancreatitis, mimicking human etiology and pathophysiology, which gave a clear answer to the long time Opie theory dilemma.

## Introduction

Acute pancreatitis (AP) is a disease with high morbidity and mortality(Banks et al., 2013). Severe acute pancreatitis, which is characterized by persistent organ failure (>48 hours), has a mortality up to 36%-50%(Buter et al., 2002; Johnson and Abu-Hilal, 2004; Mofidi et al., 2006). So far, there is no specific therapy for acute pancreatitis, and no medication has been shown to be effective in treating AP either(Banks et al., 2006; Steinberg and Tenner, 1994). Part of this reason is a lack of appropriate models.

All current models of acute pancreatitis have limitations. For example, the most popular caerulein model and its alternative models, such as the CDE diet-induced(Lampel and Kern, 1977) and the L-Arginine-induced(Dawra et al., 2007) acute pancreatitis models, have been questioned for their weak clinical relevance.

For gallstone pancreatitis, the bile salt infusion-induced mouse model of biliary pancreatitis (Laukkarinen et al., 2007) has several advantages. However, even infusion of saline can induce pancreatitis and lung injury. Another mouse model of ligation-induced gallstone pancreatitis (Samuel et al., 2010; Yuan et al., 2011) mimics the human etiology of gallstone obstruction at the biliary-pancreatic duct, but it can only produce edematous pancreatitis, which makes it not appropriate to evaluate severity of pancreatitis.

Surprisingly, both the bile salt infusion model and the ligation model use only normal mice, not gallstone formed animals. We reason that by combing formation of gallstone and ligation of the biliary-pancreatic duct, this new model could resolve questions resulted from the two old models. In this communication, we report our discoveries from the new model and its reasonable explanation of the long time dilemma of Opie theory.

## Results

### Effect of diet on severity of ligation-induced pancreatitis

To determine if gallstone formation could affect severity of acute pancreatitis, we designed three types of diet: chow, high fat and lithogenic diet. The lithogenic diet formula is 0.5% cholic acid + high fat diet. We discovered that after ligation, mice of the lithogenic diet developed the most severe damage, when compared to the group of chow diet and high fat diet (Fig. 1.). While in the sham groups, lithogenic diet did not produce any damage in the pancreas (data not shown).

**Fig 1.**
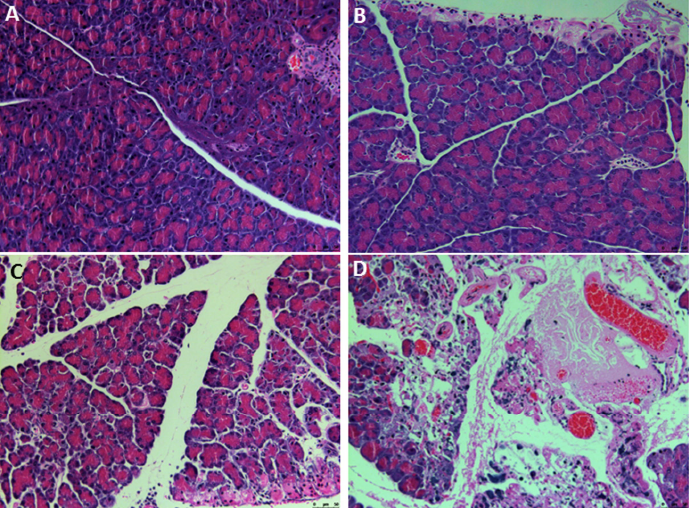
Photomicrographs of pancreatic samples obtained 24 hours after ligation of the biliary-pancreatic duct (PD). Representative samples were obtained from sham-operated mice (image A, S) and from ligation of the biliary-pancreatic duct of mice fed with chow diet (image B, PDC), high fat/cholesterol diet (image C, PDHF) and lithogenic diet (image D, PDG). Note evidence of pancreatic edema, inflammation, and massive lobular injury/ necrosis in pancreata taken from lithogenic diet fed mice (D), and mild necrosis in pancreata taken from either chow diet fed mice (B) or high fat diet fed mice(C).

We then analyzed the severity of pancreatitis using a semi-quantitative score system. Edema, inflammation and necrosis were all the most severe in the lithogenic diet group. The percentage of necrosis area to pancreas parenchyma showed a more accurate and impressive damage caused by the lithogenic diet: a 4.57-fold increase than the high fat diet group (Fig. 2.). Interestingly, in the lithogenic diet group, we didn’t see any significant difference when comparing the damage of gallstone formed mice to that of no stone (data not shown). This indicated that cholesterol supersaturated bile resulted from the lithogenic diet might play an important role in cell injury of biliary pancreatitis.

**Fig 2.**
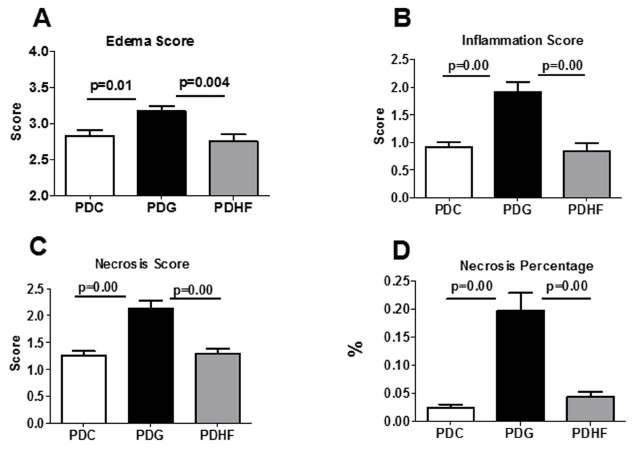
Severity of pancreatitis 24 hours after ligation of the biliary-pancreatic duct. Results were presented as mean ±SEM values obtained from 12 or more animals in each group from 3 independent experiments. A: Edema score; B: Inflammation score; C: Necrosis score; D: Necrosis percentage of pancreas parenchyma. Mice fed with lithogenic diet showed the highest damage. Methods of evaluating severity of pancreatitis was described in the text.

### Characterization of pancreatic damage distribution

We then determined the distribution characteristic of pancreas damage in the lithogenic diet group. It was thus revealed that massive cell necrosis was located mainly in the pancreas head (around 5mm of the ligation area). The pancreas tail showed almost normal tissue structure (Fig. 3.).

**Fig 3.**
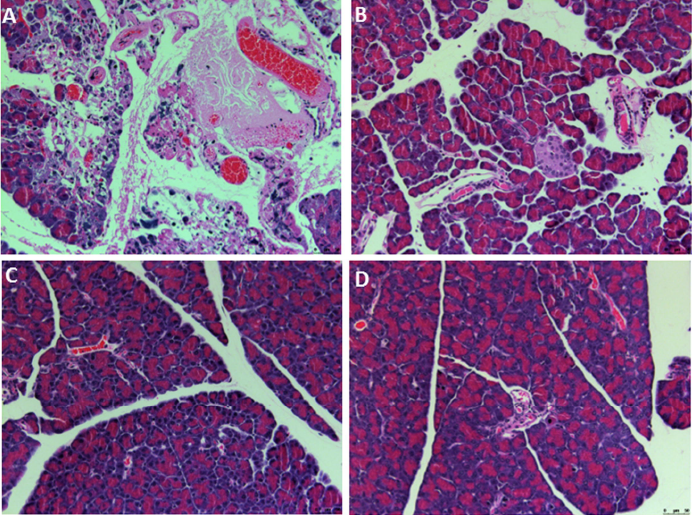
Photomicrographs of different position of pancreatic samples obtained 24 hours after ligation of the biliary-pancreatic duct from mice fed with lithogenic diet. The head area of the pancreas had the most severe tissue damage. A: head; B: neck; C: body; D: tail.

### Multi organ dysfunction of ligation-induced pancreatitis

Multiple organ dysfunction syndrome is the leading cause of severe acute pancreatitis. To determine the systemic effects on mice fed different diet, we detected organs of the lung, liver and kidney. After ligation, neutrophils infiltrated into the lung, leading to lung inflammation and injury. For each diet group, ligation always resulted in a more severe neutrophil infiltration, and the lithogenic diet group developed the most severe lung inflammation. For kidney injury represented as serum creatinine level, the lithogenic group was the most severe group again. Lastly, liver injury represented as serum AST level, was once again the most severe in the lithogenic diet group (Fig. 4.).

**Fig 4.**
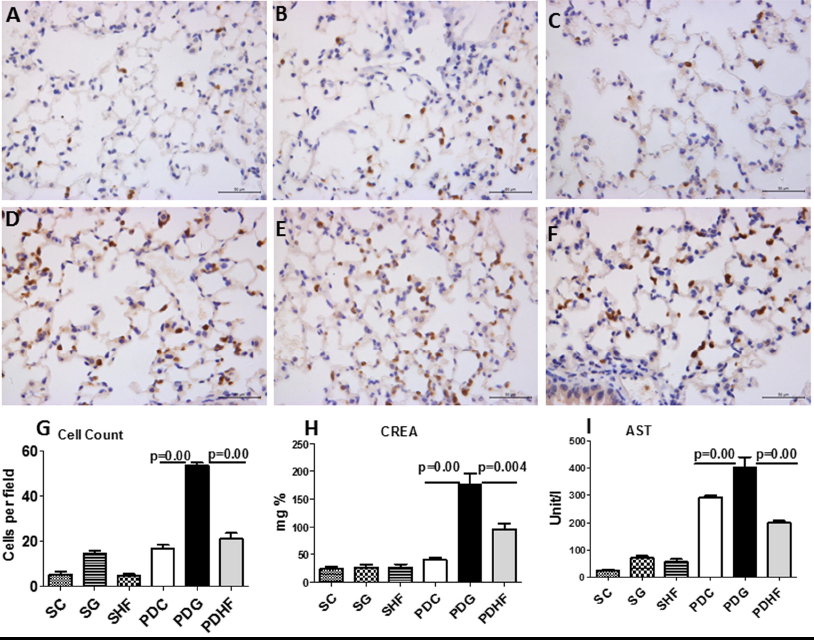
Systemic effects of the new model. Upper two panels: representative photomicrographs (Ly6G staining) of mouse lung following 24 hours of sham operation (S) or PD ligation. A: S + Chow diet (SC); B: S + Lithogenic diet (SG); C: S + high fat diet (SHF); D: PD + Chow diet (PDC); E: PD + Lithogenic diet (PDG); F: PD + high fat diet (PDHF). Graphs are described from left to right. Data are means ±SEM. Lower panel: pulmonary neutrophil count (G); plasma creatinine (H) and AST levels (I).

### Associated factors affecting severity of ligation-induced pancreatitis

As obesity has been intensively showed to increase severity of acute pancreatitis, we compared the body weight of each group. We found that the high fat diet group, instead of the lithogenic diet group, had the greatest body weight and a comparable pancreatic damage to the chow diet group, which indicates that obesity may not contribute the aggravating effects of lithogenic diet. We then measured bile components of bile of gallbladder from mice of sham group, and found that basal level of cholesterol was the highest in the lithogenic group, while level of phospholipid and bile acid didn’t show a significant change. After ligation, bile level of cholesterol and bile acid remained the highest in the lithogenic group. Bile phospholipid level was the highest in the high fat diet group (Fig. 5.).

**Fig 5.**
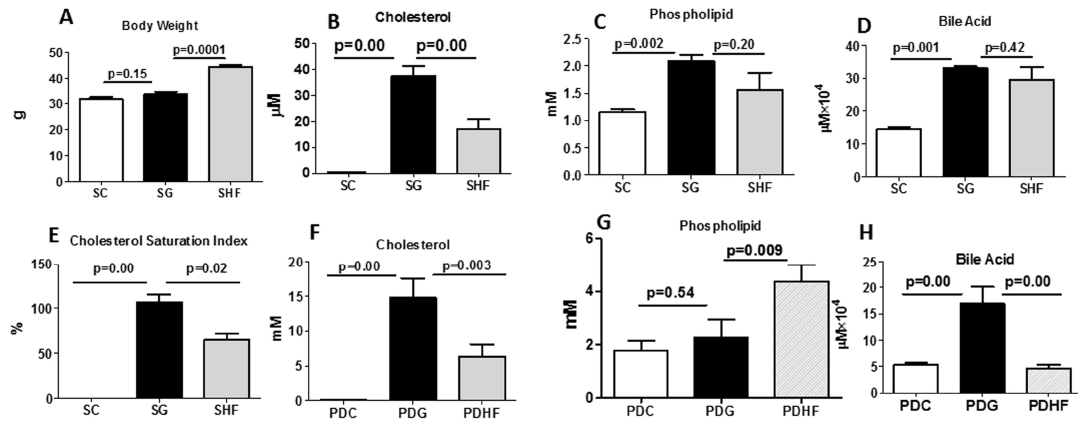
Associated affecting factors of severity of pancreatitis. Data are means ±SEM. A: body weight; B: bile cholesterol levels in sham group; C: bile phospholipid levels in sham group; D: bile acid levels in bile in sham group; E: Bile cholesterol saturation index in sham group; F: bile cholesterol levels in ligation group; G: bile phospholipid levels in ligation group; H: bile acid levels in ligation group.

### Effects of bile flow on severity of ligation-induced pancreatitis

Using the combined ligation of the common bile duct and pancreatic duct (BPD), we completedly blocked flow of supersaturated bile into the pancreas in the lithogenic diet group. The BPD group had comparable edema and inflammation to the PD group. Impressively, pancreas necrosis decreased dramatically. It decreased 90% of that PD group (PDG), which is similar to the basal level of the chow diet group (PDC, Fig. 6.).

**Fig 6.**
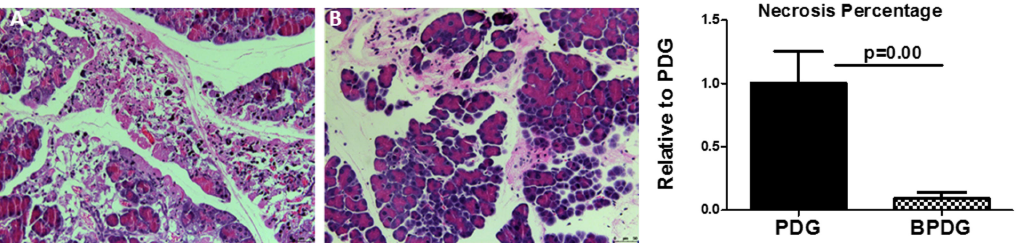
Effects of supersaturated bile on severity of pancreatitis. Once supersaturated bile was blocked from flowing into the pancreas, severity of pancreatitis was dramatically decreased. A: PD; B: BPD; C: necrosis percentage to pancreas parenchyma.

### Effects of cholesterol crystal and bile acid on viability of acinar cell

We used cellular ATP level to assess cell viability. As shown in Fig. 7, taurocholate not surprisingly caused significant cell death while cholesterol crystal alone didn’t resulted in any cell injury. Interestingly, when cholesterol crystal and taurocholate was combined together, viability of acinar cell was significantly lower than that of taurocholate alone, which indicates cholesterol crystal aggravated cell injury caused by bile acid.

**Fig 7.**
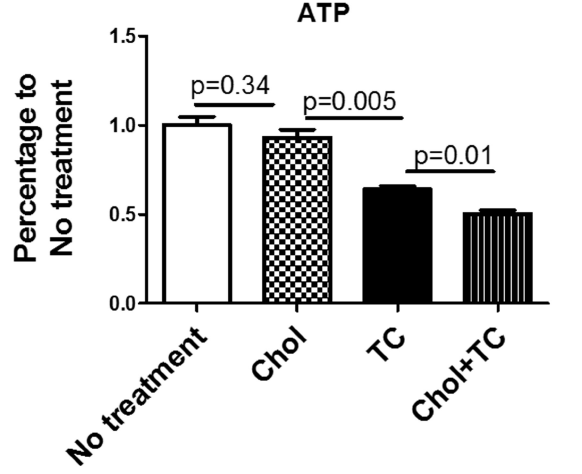
Effects of bile components on acinar cell injury. Cell injury in acini exposed to 0.5mM Na-taurocholate and/or 1 mg/ml cholesterol crystal. Acini ATP levels were evaluated as described in the text. Results shown represent mean ±SEM from three experiments performed in duplicate.

### Alteration of bile lipids metabolism pathway

Since high fat/cholesterol diet alone didn’t significantly aggravated the severity of ligation-induced pancreatitis over the chow diet, in the part of molecular mechanism, we only tested liver of the lithogenic diet group and the chow diet group. To determine whether bile lipids synthesis or transport account for the persistent high bile level of cholesterol and bile acid after ligation of the biliary-pancreatic duct, we tested several representative molecules in pathways of cholesterol/bile acid. We observed that increased levels of bile cholesterol was mainly due to up-regulation of ABCG8, a cholesterol transporter, instead of cholesterol de novo synthesis. The increased levels of bile acid was mainly from up-regulation of Cyp7a1, a rate limiting enzyme for bile acid de novo synthesis, but not from bile salt export pump (Bsep), which transport bile salt from blood in the biliary tract system (Fig. 8.).

**Fig 8.**
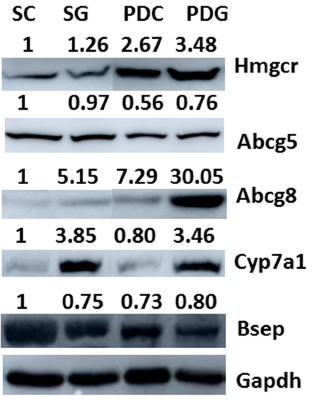
Alteration of bile acid metabolism pathway. Immunoblots of liver samples were obtained 24 hours after ligation. Densitometry ratios normalized to loading controls (GAPDH) are provided above each immunoblot lane with the sham control value represented as 1. Abbreviation was the same as described above.

## Discussion

Gallstone is the most common cause of acute pancreatitis(Forsmark et al., 2016; Lankisch et al., 2015; Yadav and Lowenfels, 2013) worldwide. Its mechanism has been recognized as obstruction caused by its passage through the terminal biliary-pancreatic duct. Unfortunately, except the opossum model(Lerch et al., 1992; Lerch et al., 1993), necrotic pancreatitis is not reproducible in other animal models of ligation-induced pancreatitis.

Based on his discoveries in autopsy, Dr. Opie proposed another mechanism, the well-known common channel or bile reflux theory(Opie, 1901a; Opie, 1901b). Though never conclusively proved, a bile infusion model was constructed based on this theory and its mouse model (Laukkarinen et al., 2007) has provided a good toll to facilitate our understanding of new mechanism.

Interestingly, both the ligation and bile salt infusion model use only normal animals, neglected formation of gallstone, the essential factor of biliary pancreatitis.

In the current study, we demonstrated that by combing formation of gallstone and ligation of the biliary-pancreatic duct, a model of necrotic pancreatitis complicated with more severe multi organ dysfunction syndrome was constructed. Compared to the ligation in the high fat/cholesterol diet group, the lithogenic diet group developed 4.6 times more necrosis area, and significantly more severe injury in the lung, kidney and liver.

Cholesterol supersaturated bile is the prerequisite for formation of gallstone(Carey, 1978; Carey and Small, 1978). In the present study, we found that after ligation of the biliary-pancreatic duct, compared to the chow diet group and the high fat/cholesterol diet group, bile levels of cholesterol and bile acid remained significantly high in the lithogenic group. When cholesterol supersaturated bile was blocked flowing into the pancreas, pancreatic necrosis area reduced to that of chow diet group. This fact indicates a role of bile components in pancreatic injury. Indeed, in the in vitro study, cholesterol crystal aggravated damage of acini caused by taurocholate.

To our knowledge, this is the first report that cholesterol crystal, the precusor of gallstone in bile, plays an important role in acute pancreatitis. In the cardiovascular system, cholesterol crystal (Duewell et al., 2010; Rajamaki et al., 2010; Samstad et al., 2014) has been reported to trigger inflammation via activating inflammasomes. While in acute pancreatitis, a recent study has reported that simvastatin is associated with reduced risk. This study is concluded from a cohort of more than three millions patient(Wu et al., 2015), which indicates an important role of cholesterol. It is worthwile investigating how cholesterol crystal acts in aggravating acute pancreatitis.

The bile reflux theory proposed by Dr. Opie is a long time controversy. Based on our data, we think that both obstruction of the pancreatic duct and reflux of bile are necessary to induce gallstone pancreatitis. If normal bile is refluxed, interstitial pancreatitis is developed. However, if cholesterol supersaturated bile is refluxed, together with changed liver function due to gallstone formation after obstrction, which maintains a persistent high bile level of cholesterol and bile acid, eventually leads to massive necrosis and worsen dysfunction of remote organs (MODS). A recent report also confirmed that the combination of bile reflux into the pancreas and obstruction of the biliary pancreatic duct could cause a more severe, necrotizing pancreatitis(Le et al., 2015). Our current study further demonstrated the important role of gallstone formation.

Our results also reveals some promising therapeutical directions in clinical practice of gallstone pancreatitis. Currently, for biliary pancreatitis patients, the only additional recommendation is to remove obstruction of the biliary pancreatic duct(Working Group, 2013). In the future, we need at least to reduce bile levels of cholesterol and bile acid for the benefits of patients, as this is a disease of two organs: liver and pancreas.

Comparing with the bile infusion model, our current model has several advantages:1, it is more severe in both local pancreas damage and remote organ dysfunction; 2, it has the pathophysiological alteration caused by gallstone; 3, it is not self-cured and suitable for evaluating therapeutical investigatios(Samuel et al., 2010; Yuan et al., 2011); 4, it is less invasive and relatively easy to operate. Of note, the bile infusion model is still important for mechanism study. For example, by infusing artificial bile with different concentration of bile components, we can demonstrate how those bile components affect severity of acute pancreatitis.

Based on our results, we proposes a novel mechanism for biliary pancreatitis: on the basis of obstruction of pancreatic duct, persistent high bile level of cholesterol and bile acid initiates and promotes acute pancreatitis.

Taken together, we generated a novel model of biliary pancreatitis which contains characteristics of both gallstone and pancreatitis. This model develops severe damage in both local pancreas and remote organs. It also provided a sound explanation for the Opie theory and some promising therapeutical directions for future clinical practice.

We expect that future studies will expand our understanding of mechanisms regarding how formation of gallstone affects the initiation and progression of biliary pancreatitis, as the site of this disease is at both the pancreas and the biliary tract system, not the pancreas only.

## Materials and Methods

### Animals

All experiments were performed using protocols approved by the Animal Care and Use Committee of Shanghai Jiaotong University. Male, 4-6 weeks age, wild-type C57BL/6 mice were purchased from Shanghai Laboratory Animal Center, Chinese Academy of Science (SLACCAS). The animals were housed in temperature-controlled (23±2°C) rooms with a 12:12-hour light/dark cycle, given freely available water.

### Induction of pancreatitis

Mice were fed the chow diet or were switched as indicated to a high fat diet (chow diet supplemented 1.25% cholesterol, 15% total fat), or a lithogenic diet (high fat diet plus 0.5% cholic acid) for 12 weeks.

Laparotomy was performed as previously described(Samuel et al., 2010) with minor modification. Briefly, 1% phenobarbital was given 50mg/kg I.P to induce anesthesia. The mice were studied in 3 experimental groups: I = sham operation; II= pancreatic duct (PD) ligation alone, the distal common bile-pancreatic duct was ligated near its junction with the duodenum; III = combined BD and PD ligation (BPD), i.e. the common bile duct was ligated first to block bile flowing into the pancreas, then, the distal common bile-pancreatic duct was ligated as in group II.

Animals were killed after 24 hours of ligation. Blood was taken for collecting serum, pancreas, liver, kidney and lung were taken for histology and molecular studies. Gallbladder bile was pooled into around 10µl each for testing bile components.

### Evaluation of pancreatitis severity

The severity of pancreatitis was evaluated by two types of method. First, edema, inflammation and necrosis were scored. The score system was defined as: For edema, 0, absent; 1, focally increased between lobules; 2, diffusely increased between lobules; 3, tense acini, widely separated lobules; 4, gross lobular separation. For necrosis, 0, absent; 1, periductal destruction (Obvious edema>2); 2, focal parenchymal necrosis (<20%); 3, diffuse loss of lobules (20% to 50%); 4, severe loss of lobules (>50%). Last for inflammation, 0, absent; 1, around ductal margins; 2, in parenchyma (<50% of lobules); 3, in parenchyma (51% to 75% of lobules); 4, massive collections. After that, to gain a more accurate data of necrotic size, pancreas necrosis area was again measured using photoshop software as necrosis area percentage(Dawra et al., 2011).

Lungs were inflated with fixative prior to excision. Selected specimens were subjected to immunohistochemistry for lymphocyte antigen 6 complex locus G6D (Ly6G). Stained cells in 10 randomly selected 20x objective fields from each lung were counted as neutrophils.

### Preparation of cholesterol crystal

Cholesterol crystal was prepared as previously reported(Duewell et al., 2010). Briefly, cholesterol was purchased from Sigma (Saint Louis, MO; cat. C8667), solubilized in hot acetone and crystallized by cooling. After six cycles of recrystallization, the final crystallization was performed in the presence of 10% endotoxin-free water to obtain hydrated cholesterol crystals. No endotoxin was detected in cholesterol crystal prepared by this method.

### In-vitro studies

Dispersed mouse pancreatic acini was prepared as previously reported with minor modification (Williams et al., 1978). Briefly, The pancreatic acini were passed through a 150µm filter. The effects of Na-taurocholate on cell viability were quantitated by measuring the percentage of adenosine triphosphate (ATP) to untreated group with CellTiter-Glo Luminescent Cell Viability Assay from Promega Corporation (Madison, WI 53711, USA).

### Systemic Studies

Plasma creatinine, aspartate serum transaminase (AST) levels and bile levels of cholesterol and bile acid were measured with commercial kits (Nanjing Jiancheng Bioengineering Institute) according to manufacturer’s manuals. Western blot was used to detect liver proteins of 3-hydroxy-3-methyl-glutaryl-coenzyme A reductase (Hmgcr), ATP-binding cassette sub-family G member 5 and 8 (Abcg5/g8), cholesterol 7 alpha-hydroxylase (Cyp7a1) and bile salt export pump (Bsep), which represents function of liver de novo synthesis and transport of cholesterol and bile acid from blood in into bile respectively. Densitometry of immunoblots was measured with Image J software (http://imagej.nih.gov/ij/).

### Analysis of data

For morphometric and other studies, analysis of parametric data was made with one-way ANOVA, while nonparametric data were compared using the Kruskal-Wallis one-way ANOVA (p <0.05), with post hoc tests for confirmation of significance.

## Acknowlegement

We thank Dr. Ashok Saluja and Dr. Rajinder Dawra at the University of Minnesota (Now University of Miami) for their selfishless guidance and constructive suggestion for this project.

This study is supported by Shanghai Key Laboratory of Pancreatic Diseases, grant 030134-01.

